# Alternating Angle Milling Suppresses Streaking Artifacts in FIB-SEM Imaging

**DOI:** 10.1101/2025.09.25.678572

**Authors:** Gleb Shtengel, Wei Qiu, Jesse Aaron, Arthur S. Crowe, Alexey A. Polilov, Katerina Karkali, Christopher K.E. Bleck, Harald F. Hess

## Abstract

Focused ion beam scanning electron microscopy (FIB-SEM) (*1*–*3*) has been used in life sciences to produce large volumetric datasets with high resolution information on ultrastructure of biological organisms. 3D image acquisition is accomplished by serial removal of thin layers of material using focused ion beam (FIB) milling followed by scanning electron microscopy (SEM) imaging.

One of the challenges in the standard FIB-SEM imaging protocol is that FIB milling results in characteristic artifacts, known as “streaks” or “curtains”. These streaks are caused by non-uniform material removal forming long straight trenches parallel to the FIB milling direction. These artifacts get worse along the milling direction and ultimately limit size of the SEM field of view.

Various methods have been proposed to mitigate the streaks in acquired images. While these techniques often provide noticeable visual improvement, the underlying problem remains. The structural information in the “streaked” areas is lost due to non-uniform material removal during milling and cannot be fully recovered.

We propose a simple modification allowing for a significant reduction of milling non-uniformities of streaks. We demonstrate the effectiveness of this approach on various samples.

## Introduction

Many commercial and research FIB-SEM systems use focused beam of Ga^+^ ions to precisely ablate thin layers of material between consecutive SEM imaging steps. Exposure to the beam of electrons hardens the surface of the epoxy-embedded samples during SEM imaging. The degree of hardening depends on the local density of heavy metal staining and is often non-uniform. As a result, material removal downstream of these non-uniformities may be also non-uniform, resulting in streaking artifacts. Examples of streaks are shown in Figure 2(a, b), and Figure 3 (a, b). Because the milling rate is non-uniform in the areas of the streaks, they form grooves on the top surface of the milled sample. As a result, the ultrastructural shapes are distorted, and in the worst cases there are noticeable membrane discontinuities (Figure 2 (b)). More examples of the FIB-SEM data sets with streaking artifacts are available on OpenOrganelle data portal (*4, 5*).

Severity of streaks depends on the choice of embedding resin, Epon is generally considered more prone to streaking artifacts than Durcupan. Finally, streaks get worse with increasing electron dose during SEM imaging, setting another constraint in the parameter space of imaging conditions.

De-streaking techniques have been proposed as a part of post-processing workflow. These methods include variational (*6, 7*) and Fourier filtering (*8, 9*) approaches.

These methods generally improve the visual appearance of the images, but two problems remain. The methods based on filtering to remove the “streak” content also remove some of the actual data content, and they do not add any data in place of the removed data. Moreover, the strong streaks have three-dimensional (3D) nature as can be seen in Figure 2 (b), and 2D image processing algorithms cannot remove these 3D-image distortion artifacts.

Recently FIB spin milling serial SEM approach (*10, 11*) has been proposed, where ion flux is delivered to the sample at a near-glancing angle (1º to 4º) from several different azimuthal directions, reducing curtaining artifacts. That configuration is not compatible with systems using highly focused Ga^+^ beams and utilizing feedback for precise milling (*2, 3*), but the concept of milling from different angles to reduce milling nonuniformities is promising.

### Proposed Approach

We propose using alternating angled milling as a way of avoiding streak artifacts. A standard FIB-SEM procedure uses FIB-milling at fixed angle, as shown in Figure 1 (top).

**Figure 1.**
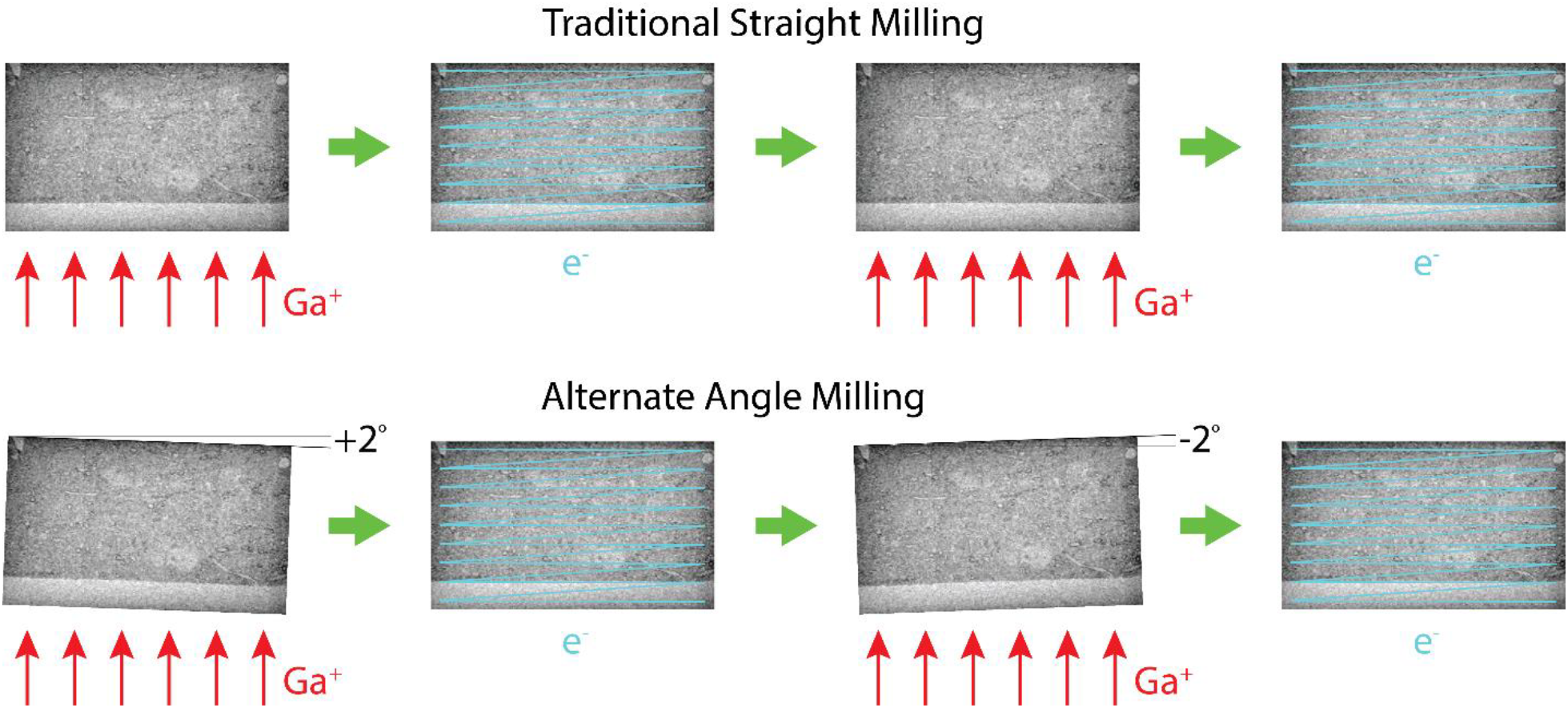
Traditional straight milling step sequence (top) and proposed alternate angle milling step sequence (bottom).

The proposed modification to the standard procedure is that the sample is rotated by a small amount (we used ±2º) for the milling step and then rotated back into original orientation for the imaging step as shown in Figure 1 (bottom). The rotation angle is inverted between subsequent milling steps, thus preventing the streak formation.

### Experimental Results

We modified the custom hardware and software for FIB-SEM described in (*2, 3*) to allow for alternate angle stage rotation between the FIB milling and SEM imaging cycles.

We also modified the sample holder configuration. It is important to minimize the lateral sample displacement during rotation so that its exposure to the FIB beam at alternate milling angles is as similar as possible. To allow for precise sample holder centering, we reduced the size of the intermediate sample holder and added shims, as illustrated in SI Figure 1.

We tested the proposed procedure on two samples with known “streak” artifacts. One sample was a *Megaphragma* wasp embedded in Durcupan (*12*), imaged with following conditions: field of view = 70 x 150 µm, pixel size = 8 nm, EHT = 1.2 kV, SEM current = 2 nA, pixel dwell time = 1µs. The second sample was *Drosophila* brain embedded in Epon (*13*), imaged with following conditions: field of view = 50 x 50 µm, pixel size = 8 nm, EHT = 1.2 kV, SEM current = 2 nA, pixel dwell time = 1µs. The pixel size, SEM current, and dwell time in both cases corresponded to the imaging dose of 195e^-^/nm^2^.

Both samples were imaged using standard protocol (straight milling) and modified protocol (alternating ±2º angled milling). The typical SEM frames for these samples are shown in Figure 2 (a, b) and Figure 3 (a, b) for straight milling and in Figure 2 (e, f) and Figure 3 (e, f) for alternating ±2º angled milling.

**Figure 2.**
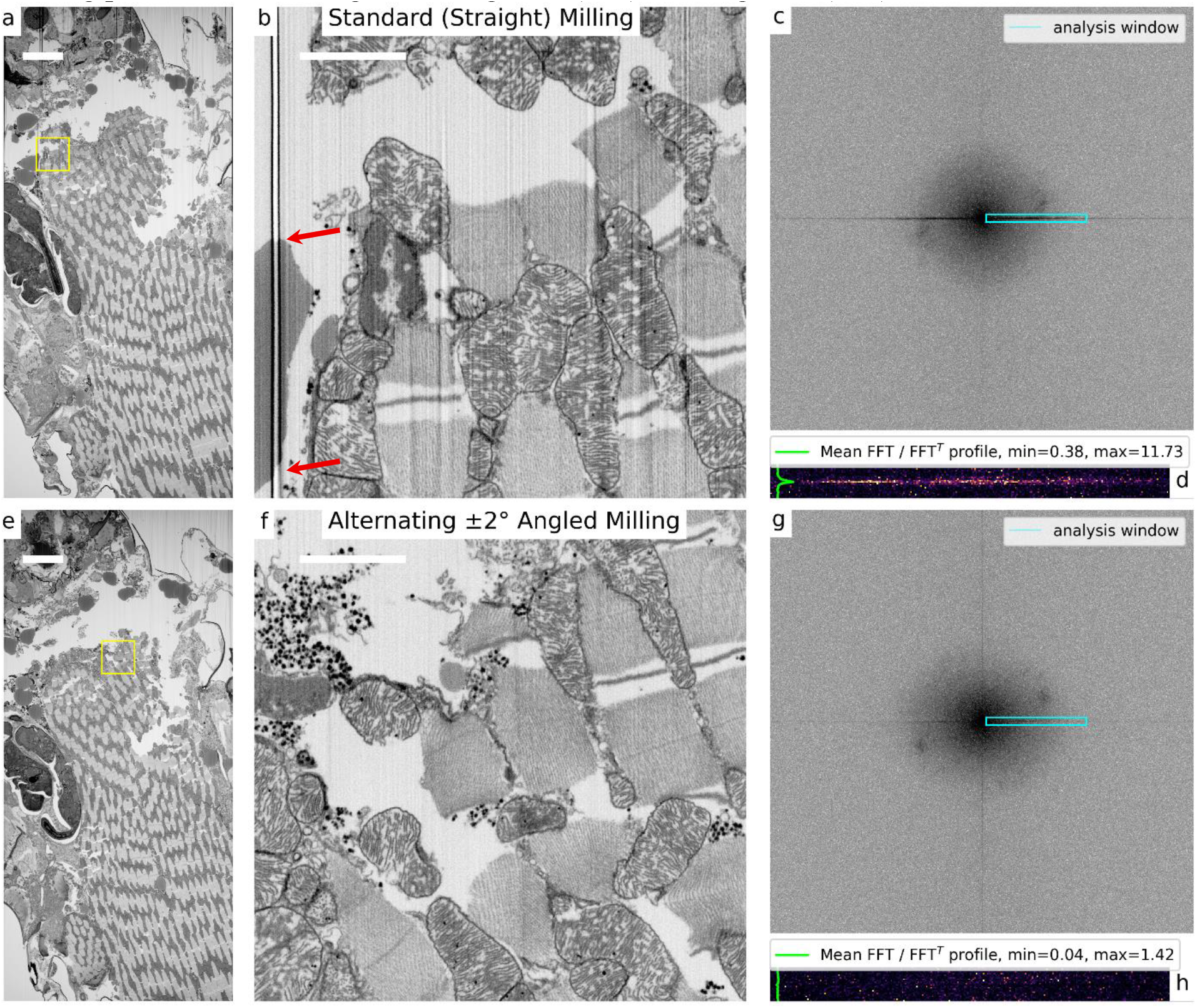
SEM images and streak analysis performed on 70x150 µm Durcupan-embedded sample of *Megaphragma* imaged under standard milling (top row) and alternating ±2º angled milling (bottom row) conditions. Original SEM frames (a, e), zoomed single tile subsets (b, f), FFT of these tiles, and streak magnitude analysis (d, h). Red arrows in (b) indicate membrane discontinuities in the area of strong streak. The scale bars are 10 µm (a, e), and 2 µm (b, f).

We collected over 900 frames under each condition for the Durcupan-embedded sample and then over 1000 frames under each condition for Epon-embedded sample.

In both cases we observed significant reduction or complete elimination of streaks with no adverse side-effects. To quantify the improvement offered by the new method, we evaluated the streaks using Fast Fourier Transform (FFT) analysis. Each acquired SEM image was split into tiles of 1250x1250 pixels (10 x 10µm). Examples of such tiles are indicated by yellow boxes in Figure 2 (a, e), and Figure 3 (a, e) and shown at larger zoom in Figure 2 (b, f), and Figure 3 (b, f). FFT was performed on each tile (Figure 2 (c, g), and Figure 3 (c, g)). As can be seen from these FFT maps, streaks have very distinct signature - increased magnitude in the areas along FFT x-axis (Figure 2 (c), and Figure 3 (c)). Dividing the FFT by the transposed FFT allows for normalization of the data, the examples of 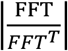 for a single frame are shown in Figure 2 (d, h), and Figure 3 (d, h). Averaging the 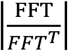 data over all frames acquired under a given condition improves the accuracy of the analysis. We then select a subset (indicated by cyan rectangle in Figure 2 (c, g), and Figure 3 (c, g)) and then average the data along FFT *x*-axis to further reduce the noise. The resulting profile is shown in green in Figure 2 (d, h), and Figure 3 (d, h).

**Figure 3.**
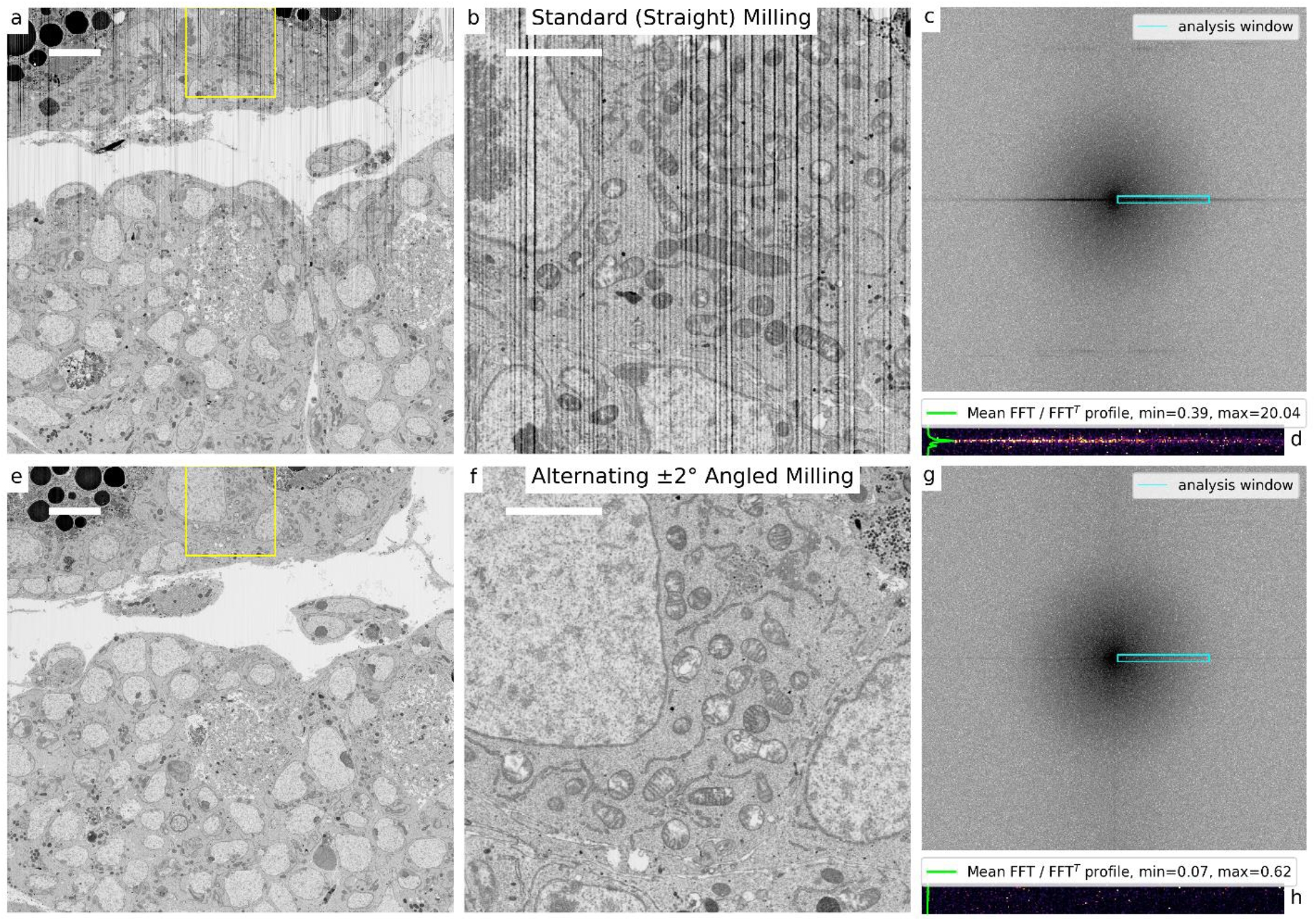
SEM image and streak analysis performed on 50x50 µm Epon-embedded sample of *Drosophila* imaged under standard milling (top row) and alternating ±2º angled milling (bottom row) conditions. Original SEM frames (a, e), zoomed single tile subsets (b, f), FFT of these tiles (c, g), and streak magnitude analysis (d, h). The scale bars are 5 µm (a, e), and 2 µm (b, f).

We define the streak magnitude as the maximum value of that profile:

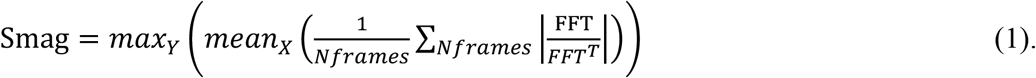

We used this metric to characterize streak magnitudes under different conditions. To increase the accuracy, we employed further averaging steps – over multiple frames and then over multiple tiles along *x*-direction, see SI for more details. The results are shown in Figure 4. It should be noted that even after averaging, there is still some noise in determining streak magnitude using the procedure outlined above, and the results are not accurate when streak magnitude value falls below 0.5. Still, we see almost complete streak suppression in both samples when the new alternating ±2º angled milling protocol.

**Figure 4.**
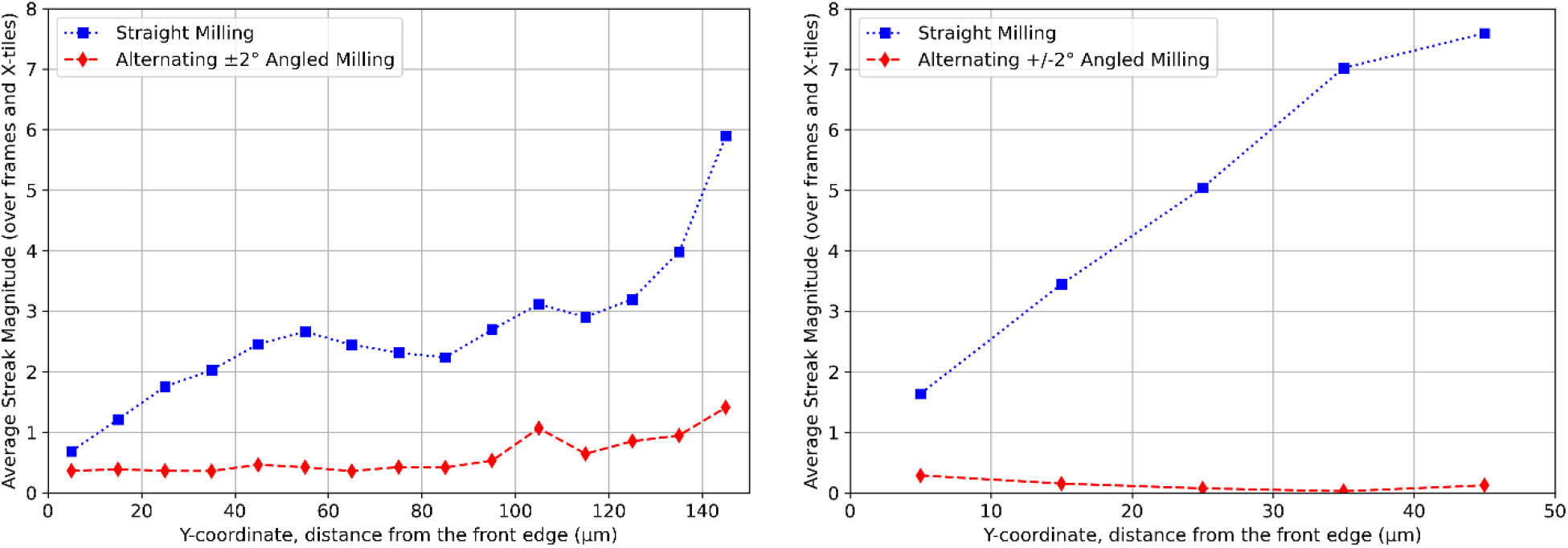
Streak magnitude vs. distance from the front edge of the FOV for *Megaphragma*, Durcupan-embedded 70x150 µm sample (left) and for *Drosophila*, Epon-embedded 50x50 µm sample (right) imaged using straight milling (blue squares) and alternating ±2º angled milling (red diamonds) protocols.

We wanted to make sure that the sample rotation under the alternate angle milling protocol does not have adverse side effects, such as reduced image quality. We have registered the data using SIFT-based approach with Affine transformation regularized to rigid translation as a transformation model (*14*) and analyzed the quality of the stack registration as well as image resolution.

It should be noted that the streaks affect the results of image quality evaluation. Sharp streaks result in increased SNR and cross-correlation values. To minimize such bias in using these metrics we selected the evaluation areas least affected by streaks, as close to the front edge of the sample as possible. Finally, the sample details may also affect the value of the metrics such as Normalized Cross-Correlation (NCC) and Signal to Noise Ratio (SNR) (*14*). We tried to select the evaluation areas to have as similar sample features as possible to reduce this uncertainty.

The results are shown in Table 1, SI Figure 5, and SI Table 1. The results are nearly identical for standard straight milling and alternating ±2º angled milling protocols.

**Table 1.** Results of evaluation of registration and Image quality.

| Evaluation Metric | *Megaphragma*, Durcupan-embedded |  | *Drosophila*, Epon-embedded |  |
| --- | --- | --- | --- | --- |
| | Straight Milling | Alternating $\pm 2^\circ$ angled milling | Straight Milling | Alternating $\pm 2^\circ$ angled milling |
| Normalized Cross-Correlation (NCC) | 0.923±0.015 | 0.920±0.014 | 0.930±0.012 | 0.928±0.017 |
| Signal to Noise Ratio (SNR) | 12.3±2.33 | 11.9±2.26 | 13.6±2.43 | 13.6±3.64 |
| Resolution (nm) | 5.05±2.49 | 5.07±2.50 | 5.30±2.62 | 5.21±2.56 |

We have also compared the image resolution in the data sets acquired using the straight milling and alternating ±2º angled milling protocols. We used 37% to 63% transition length as a resolution criterion (*3*). The details of the blob statistics are presented in SI Table 2, SI Figure 9 and SI Figure 9. The examples of the blobs and edge transition analysis procedure are shown in SI Figure 10, SI Figure 8, SI Figure 10, and SI Figure 11. The resolution analysis results are nearly identical for standard straight milling and alternating ±2º angled milling protocols.

We conclude that the proposed alternating ±2º angled milling protocol significantly reduces the streaking artifacts and does not affect the quality of the SEM imaging as well as the quality of eventual stack registration.

## Conclusions

We have proposed a simple modification of FIB-SEM acquisition protocol that utilizes alternating ±2º angled milling to significantly reduce streaking artifacts.

The proposed protocol can be utilized with no hardware redesign and requires only a minor change in acquisition software and sample holder hardware.

We confirmed significant streak suppression under new acquisition protocol, extending artifact-free depth of SEM imaging to 150 μm in Durcupan-embedded sample and to 50 μm in Epon-embedded sample.

We observed no adverse side effects to the quality of the acquired SEM data.

## Acknowledgements

We thank Ken Hayworth, David Peale, and C. Shan Xu for helpful discussions.

## Author information

### Contributions

HFH conceived the concept and oversaw the project, GS modified the software and hardware, AAP and KK provided the samples, GS, WQ, ASC, and JA performed the measurements, GS processed the data and performed the analysis, all authors discussed the results and wrote the correspondence.

## Supplemental Information

### Hardware Modification

The hardware was modified to allow for precise adjustment of the sample position so that the milled tab is located at the rotation center of the SEM stage. The sides of the intermediate SEM sample holder were milled down as shown in SI Figure 1 (right). The added metal shims allow for precise *x-y* positioning of the intermediate sample holder to make sure that the SEM sample tab is at the center of the SEM stage rotation. This was tested using SEM as shown in SI Figure 2. The SEM stage was rotated by ±2º and SEM images were acquired to confirm that these angular displacements result in minimal lateral displacements.

### Source Code

The experiment control software (written in National Instruments Labview) is available through Janelia Open Science Tools & Innovations sharing: https://www.janelia.org/open-science/enhanced-fib-sem

Data processing was performed in Python using the FIBSEM_gs_py library (*14*). Python Jupyter notebooks are available: https://github.com/gleb-shtengel/FIBSEM_gs_py/blob/main/examples/Register_FIB-SEM_stack_DASK_v4_J5_jrc_Megaphragma_2023_3n4_Angled_Milling_Analysis_Durcupan.ipynb and https://github.com/gleb-shtengel/FIBSEM_gs_py/blob/main/examples/Register_FIB-SEM_stack_DASK_v4_jrc_Blanco_MutDrosophila_Old_J5_Angled_Milling_Analysis_Epon.ipynb

### Estimation of Streak Magnitude

The streak magnitude for each tile within SEM image was defined in Eq (1) as the maximum value of the 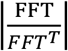 within a cyan box averaged over X-direction in Figure 2 (c, g) and Figure 3 (c, g).

We used this metric to characterize streaking under different conditions. To increase the accuracy, we employed further averaging steps – over multiple frames and then over tiles along *x*-direction.

The first step is to split each frame into tiles and calculate the absolute value of FFT for each tile, as shown in SI Figure 3.

We then calculated the streak magnitude for each tile using the Eq (1).

This procedure is performed on each frame acquired under a given condition, then the results are averaged over all frames collected for a given condition SI Figure **4** (left). Finally, the data is averaged over the tiles along *x*-coordinate (horizontal direction in SI Figure **4** (left)). As a result, we obtain the dependence of streak magnitude on Y-coordinate SI Figure **4** (right), and data in Figure 4.

### Estimation of the Registration Quality

Image stacks were registered using SIFT-based approach with Affine transformation regularized to rigid translation as a transformation model (*14*). We then determined Normalized Cross Correlation (NCC) for sequential frames within each stack. NCC vs. Frame for both samples are presented in SI Figure 5. We also determined Signal to Noise Ratio (SNR) from NCC using the following equation (*15*):

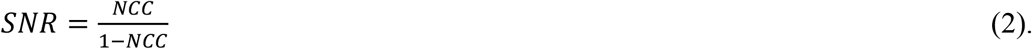

The results are summarized in SI Figure 5 SI Table 1. The results are nearly identical for standard straight milling and alternating ±2º angled milling protocols.

### Estimation of the Image Resolution

Image resolution was determined using the procedure described in (*3*). We used skimage implementation of Laplacian of Gaussian (LoG) (*16*) for initial blob detection. For each blob we analyzed the rising and falling edges along X- and Y-directions. We used 37% to 63% transition length as a resolution criterion. The details of the blob statistics are presented in SI Table 2, SI Figure 9 and SI Figure 9. The examples of the blobs and edge transition analysis procedure are shown in SI Figure 10, SI Figure 8, SI Figure 10, and SI Figure 11. The resolution analysis results are nearly identical for standard straight milling and alternating ±2º angled milling protocols.

---

**SI Figure 1.**
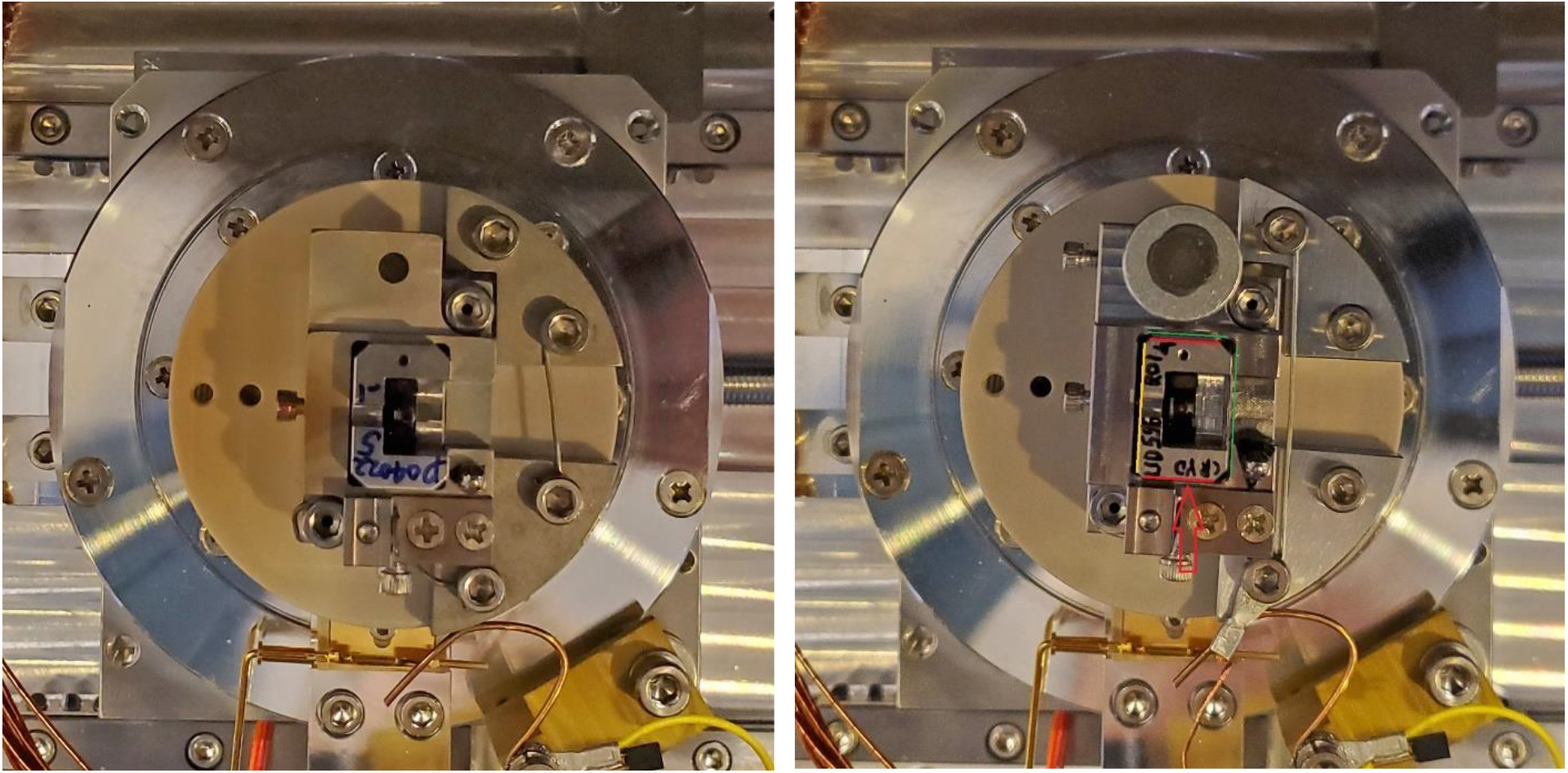
Modification of the intermediate sample holder and sample holder assembly. Original sample holder arrangement is shown in the left photo, and modified arrangement in the right photo. The size of the intermediate sample holder (indicated by the red arrow in the right photo) was reduced. The back side (indicated by the yellow line in the right photo) was milled by 0.50mm, the sides (indicated by the red lines in the right photo) were milled down by 0.25mm each. The shims were added along the walls of the outside sample holder indicated by green lines in the right photo.

**SI Figure 2.**
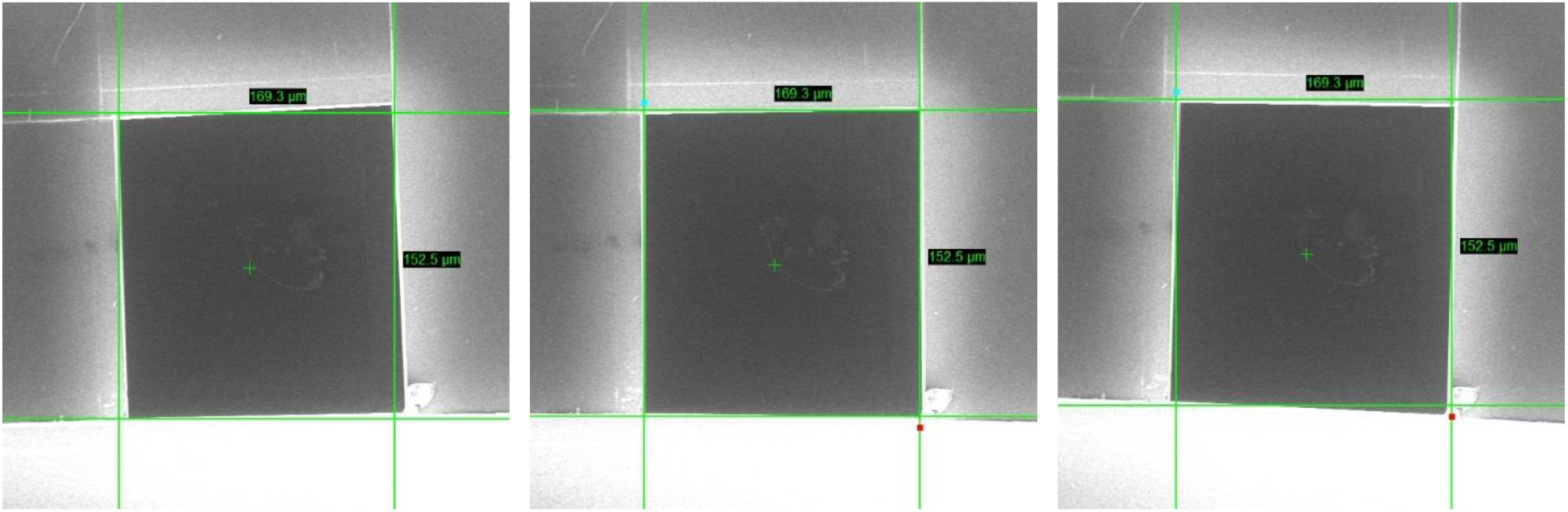
Images of the SEM sample at different SEM rotation stage positions. SEM imaging is done at SEM rotation stage position 194.5º (center). Milling is done at ±2º displacements, the stage is rotated to 192.5º (left) and to 196.5º (right). If the sample is positioned so that the SEM sample tab is at the center of the stage rotation, these angular displacements will result in minimal lateral displacements.

**SI Figure 3.**
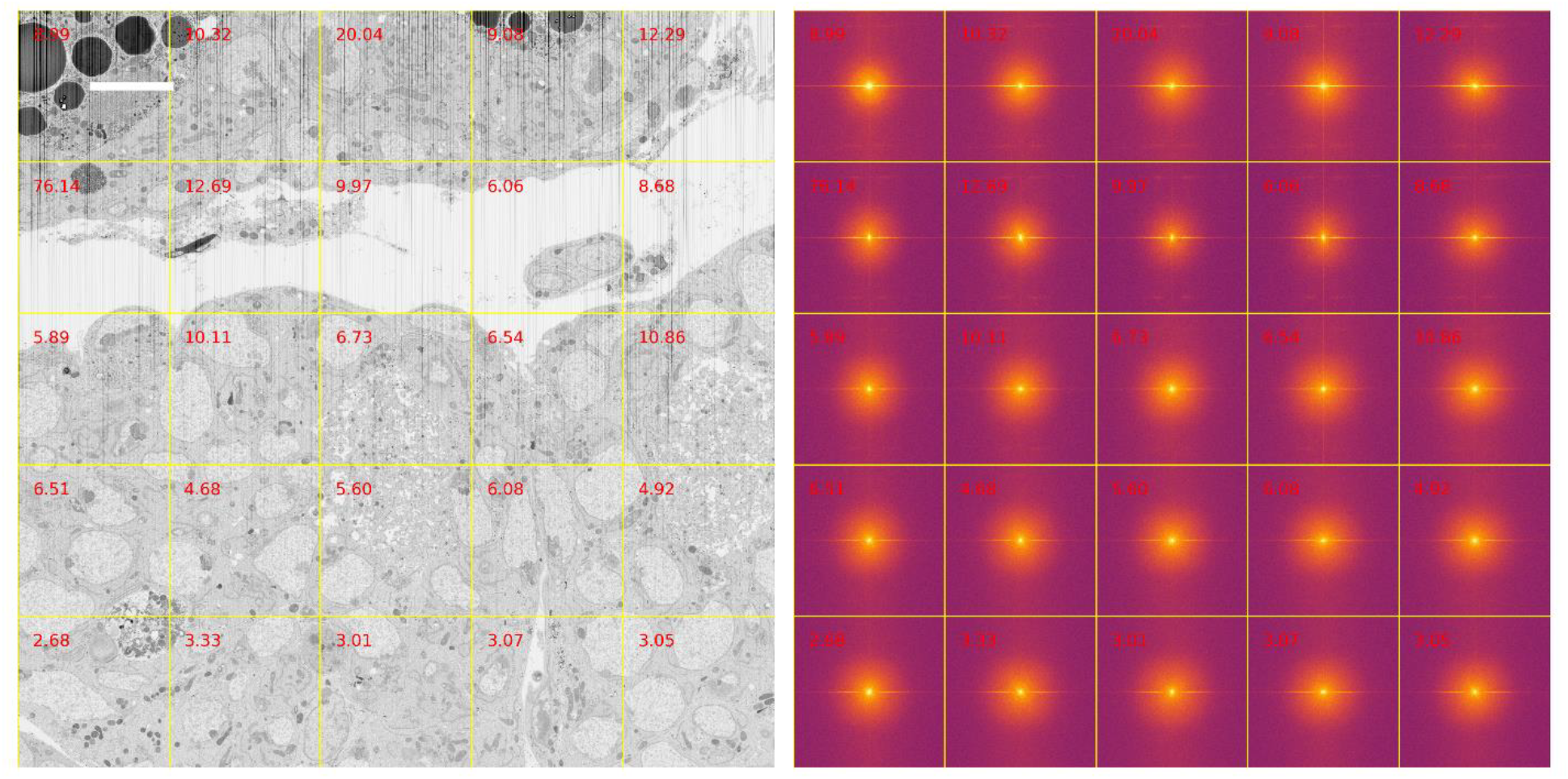
Single SEM frame (*Drosophila*, Epon-embedded sample, straight milling conditions) split into tiles (left). FFT is calculated for each tile (right). Scale bar is 5 µm

**SI Figure 4.**
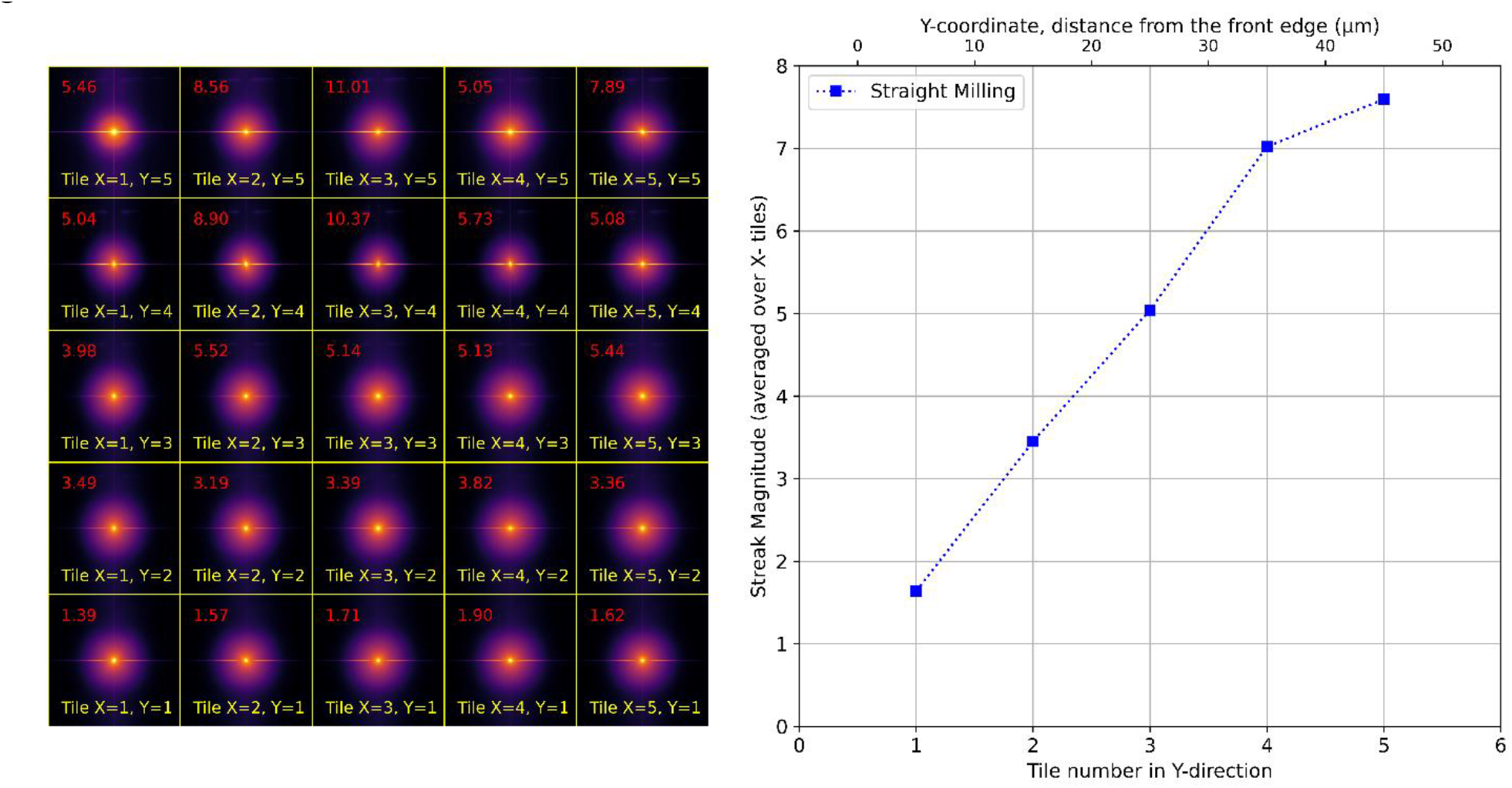
Tiled FFT averaged over all frames acquired for *Drosophila*, Epon-embedded sample under straight milling protocol (left). Tile numbers in X- and Y-direction are in yellow, streak magnitude for each tile is in red. Streak magnitude averaged over tiles in X-direction vs Tile number in Y-direction (right).

**SI Figure 5.**
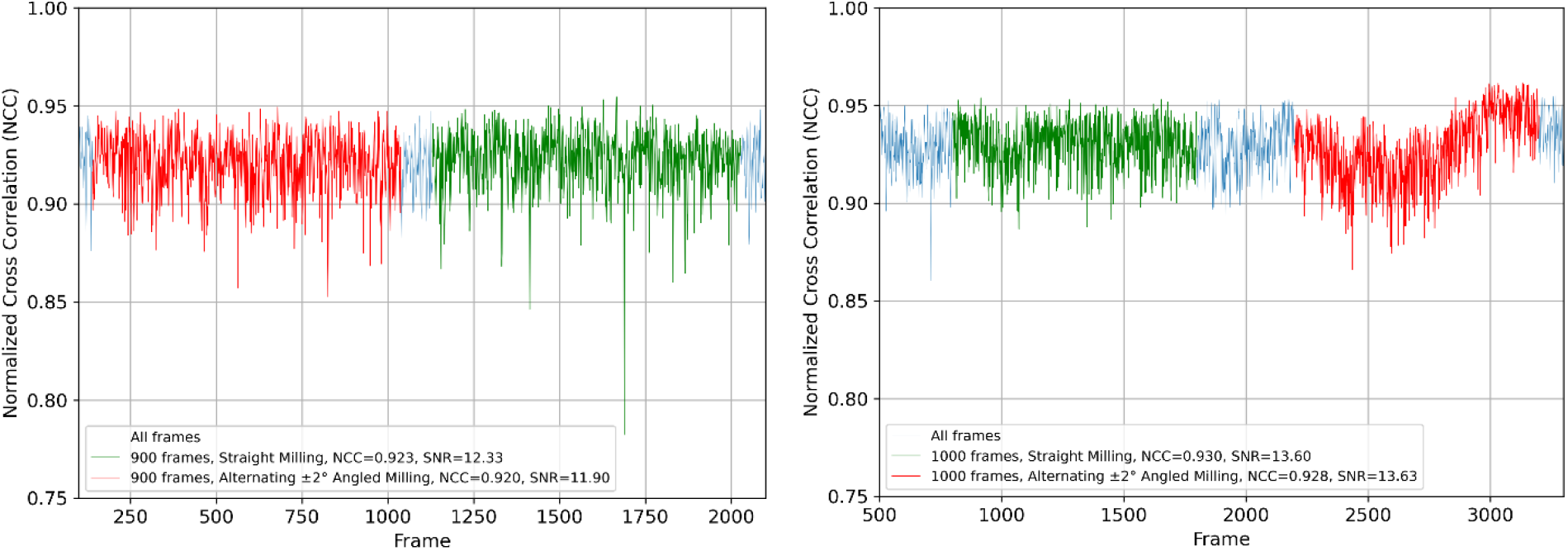
Normalized Cross Correlation vs. Frame for the samples *Megaphragma*, Durcupan-embedded sample (left) and for *Drosophila*, Epon-embedded sample (right) imaged using straight milling (green section of the trace) and alternating ±2º angled milling (red section of the trace).

**SI Figure 6.**
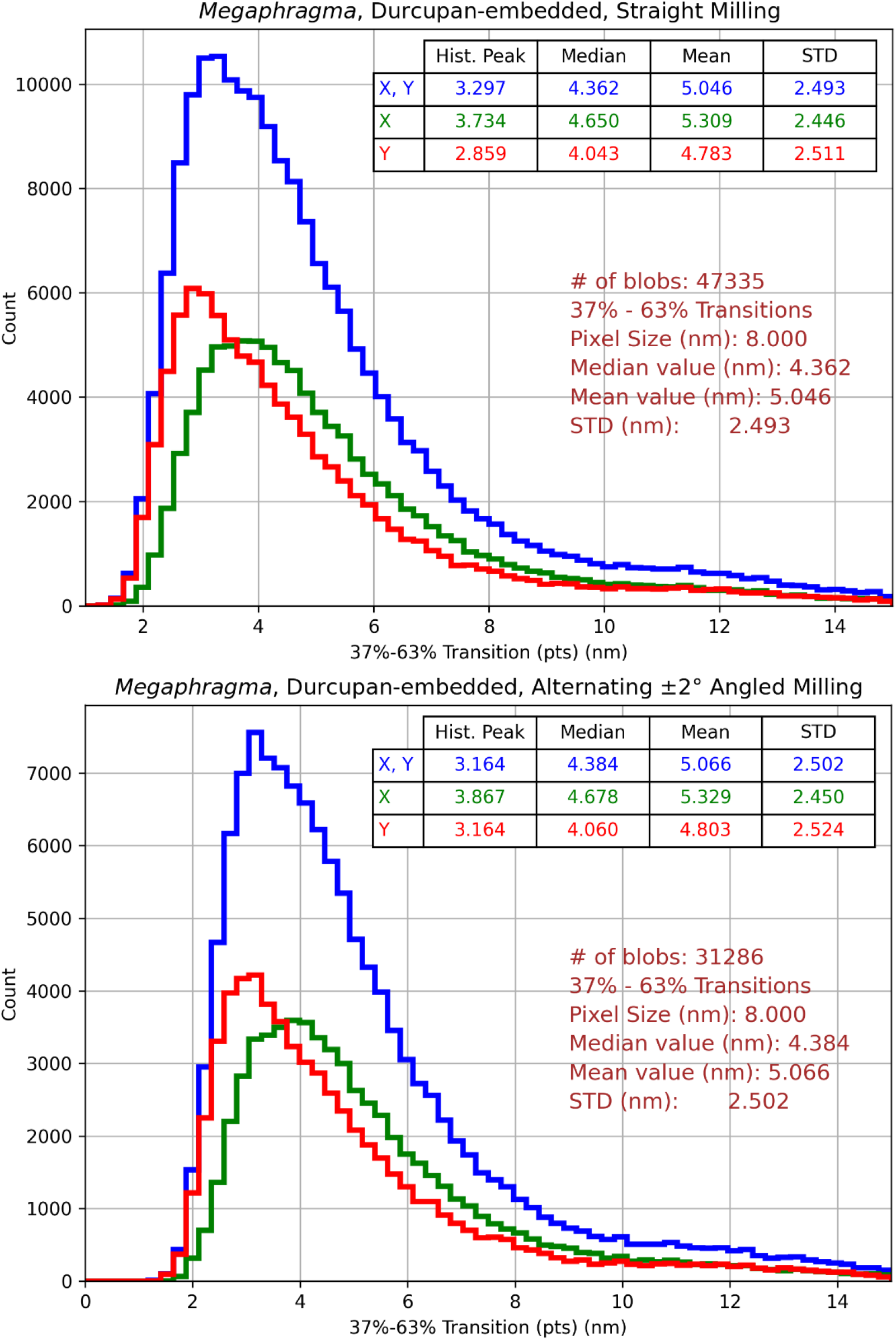
Distributions of X- and Y-edge transitions in the FIB-SEM images collected on the *Megaphragma*, Durcupan-embedded sample under straight milling (top) and alternating ±2º angled milling (bottom) protocols.

**SI Figure 7.**
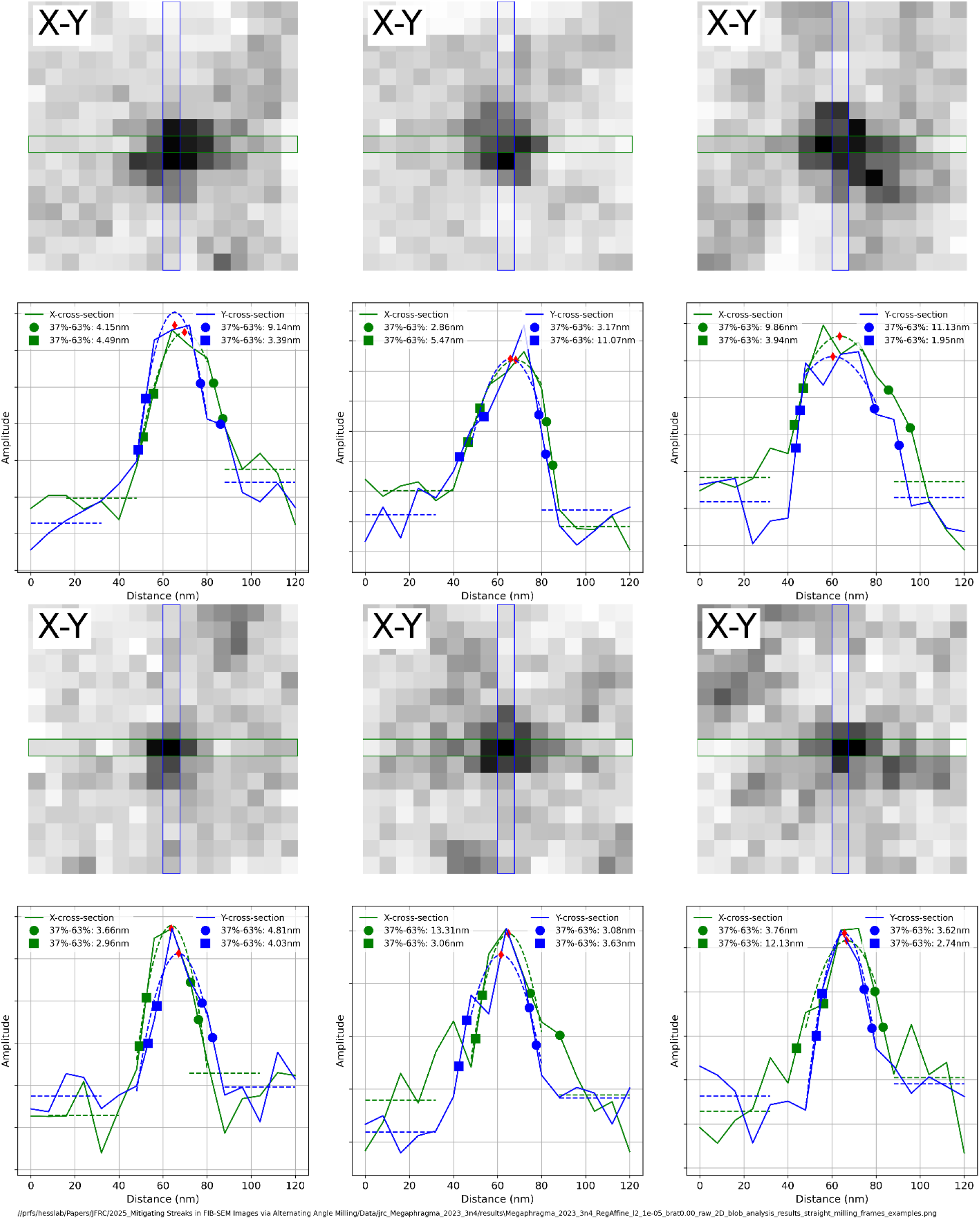
Examples of blobs used for edge transition analysis on the *Megaphragma*, Durcupan-embedded sample imaged using straight milling protocol.

**SI Figure 8.**
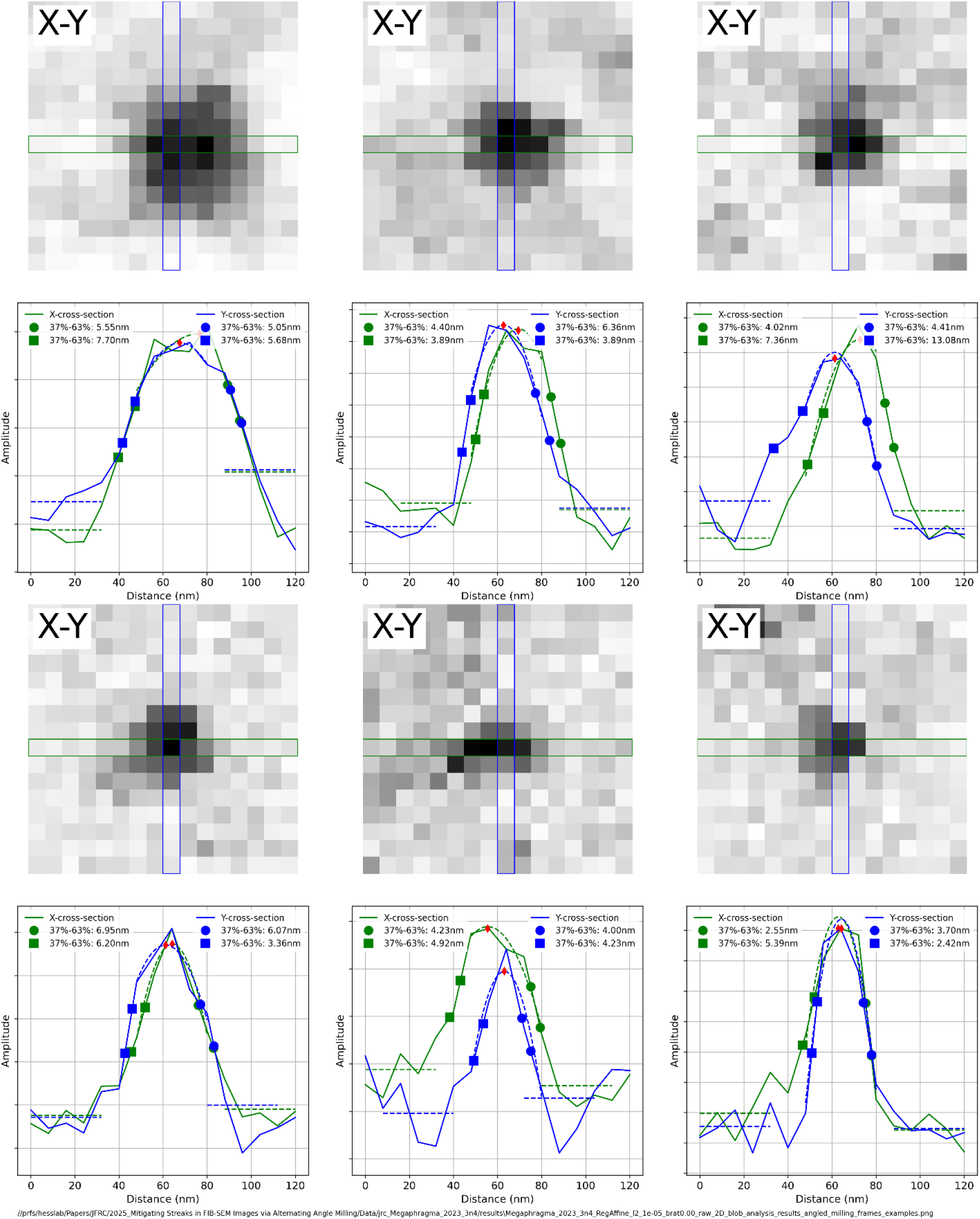
Examples of blobs used for edge transition analysis on the *Megaphragma*, Durcupan-embedded sample imaged using alternating ±2º angled milling protocol.

**SI Figure 9.**
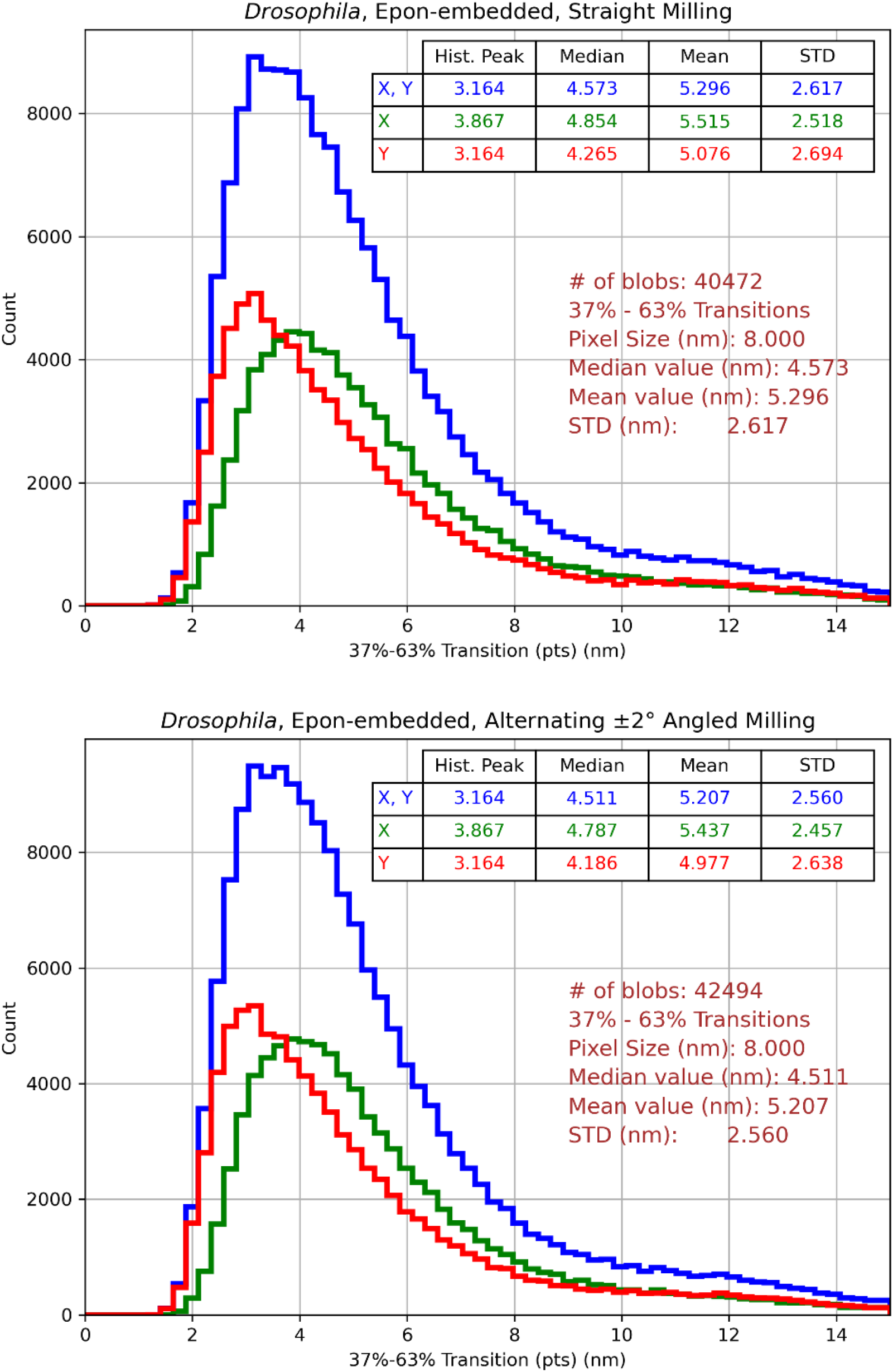
Distributions of X- and Y-edge transitions in the FIB-SEM images collected on the *Drosophila*, Epon-embedded sample under straight milling (top) and alternating ±2º angled milling (bottom) protocols.

**SI Figure 10.**
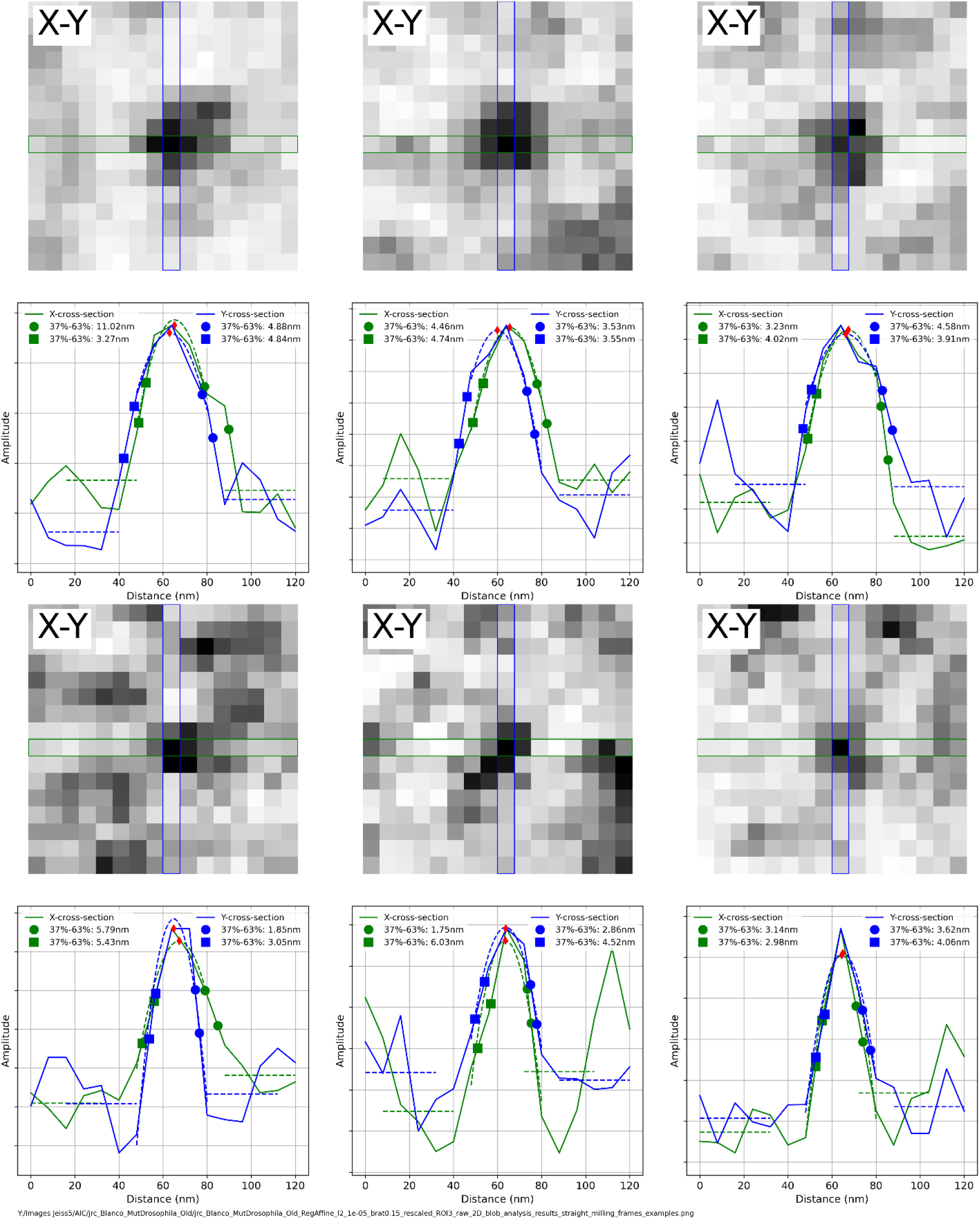
Examples of blobs used for edge transition analysis on the *Drosophila*, Epon-embedded sample imaged using straight milling protocol.

**SI Figure 11.**
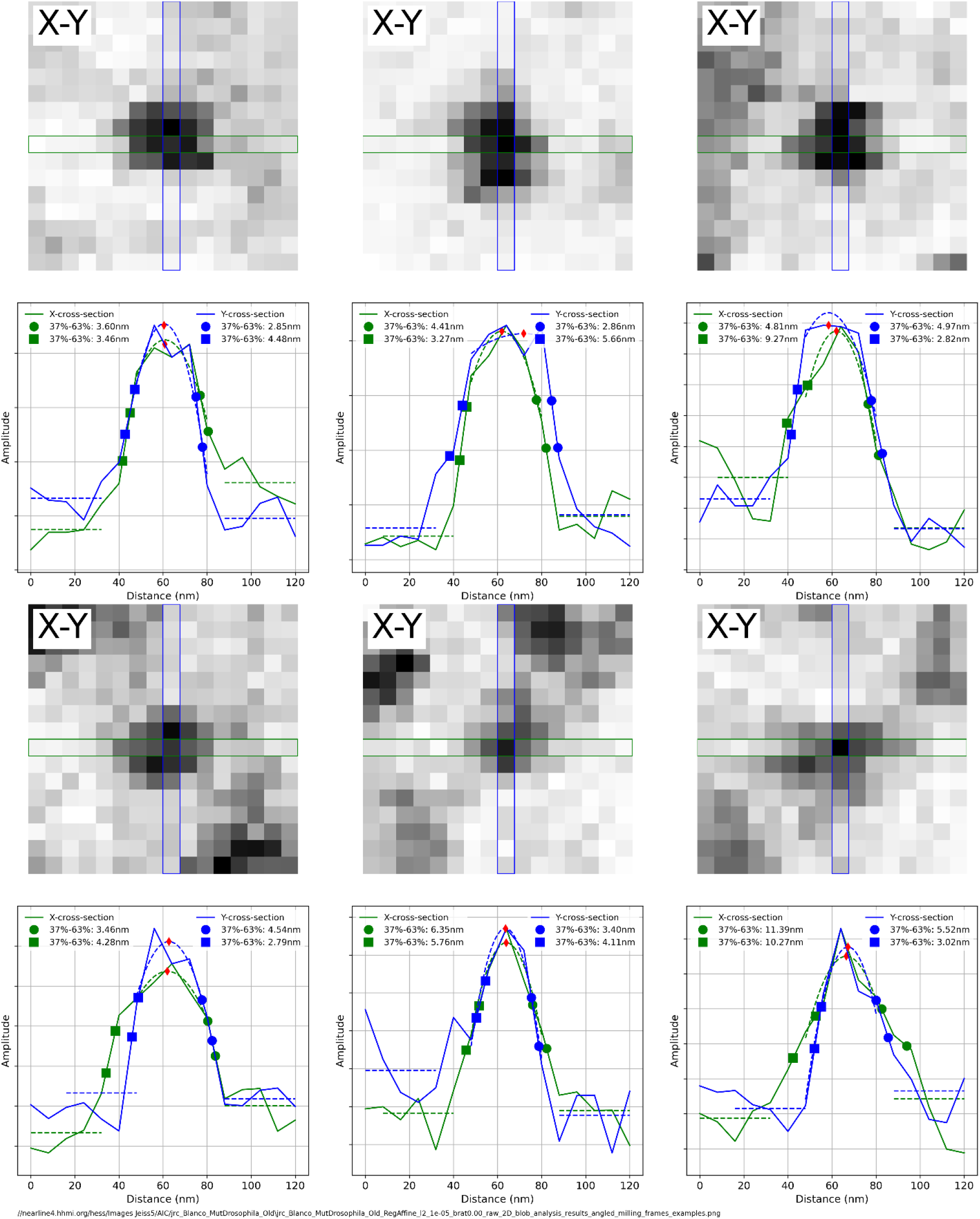
Examples of blobs used for edge transition analysis on the *Drosophila*, Epon-embedded sample imaged using alternating ±2º angled milling protocol.

**SI Table 1.** Results of evaluation of registration and Image quality.

| Evaluation Metric | *Megaphragma*, Durcupan-embedded |  | *Drosophila*, Epon-embedded |  |
| --- | --- | --- | --- | --- |
| | Straight Milling | Alternating $\pm 2^\circ$ angled milling | Straight Milling | Alternating $\pm 2^\circ$ angled milling |
| Number of frames used for analysis | 900 | 900 | 1000 | 1000 |
| Normalized Cross-Correlation (NCC) | 0.923±0.015 | 0.920±0.014 | 0.930±0.012 | 0.928±0.017 |
| Signal to Noise Ratio (SNR) | 12.3±2.33 | 11.9±2.26 | 13.6±2.43 | 13.6±3.64 |

**SI Table 2.** Details of the estimation of the image resolution.

| Evaluation Metric | *Megaphragma*, Durcupan-embedded |  | *Drosophila*, Epon-embedded |  |
| --- | --- | --- | --- | --- |
| | Straight Milling | Alternating $\pm 2^\circ$ angled milling | Straight Milling | Alternating $\pm 2^\circ$ angled milling |
| Number of frames used for analysis | 900 | 900 | 1000 | 1000 |
| Number of detected blobs | 47335 | 31286 | 40472 | 42494 |
| Resolution (nm) | 5.05±2.49 | 5.07±2.50 | 5.30±2.62 | 5.21±2.56 |

